# An Improved Linear Mixed Model for Multivariate Genome-Wide Association Studies

**DOI:** 10.1101/2022.02.21.481252

**Authors:** Dan Wang, Jun Teng, Changheng Zhao, Xinhao Zhang, Hui Tang, Xinzhong Fan, Shizhong Xu, Qin Zhang, Chao Ning

## Abstract

Current methods of multivariate analysis require complete multivariate phenotypes from each individual and have a computational time complexity of *O*(*n*^2^) per SNP, where *n* is the sample size. We develop an efficient genomic multivariate analysis tool (GMAT) for genome-wide association studies of multiple correlated traits. The new method can handle incomplete multivariate data with missing records and reduce the time complexity to *O*(*n*) per SNP. Simulation studies based on known genotypes and phenotypes of actual populations show that GMAT has increased the statistical power with a proper control of false positivity for association studies compared to the conventional linear mixed model (LMM) that removes individuals with incomplete records. Applications to a balanced donkey data and an unbalanced yeast data show that the computational efficiency of the new method has been increased about tens of times faster than the conventional LMM analysis. The GMAT package can be downloaded at https://github.com/chaoning/GMAT.

## 1 Background

Genome-wide association studies (GWAS) are becoming valuable tools for investigation of the genetic architecture of economically important quantitative traits. Over the last two decades of practice of GWAS, many replicated genomic loci have been discovered to be associated with complex diseases and quantitative traits [1]. The common practice of GWAS is to identify the association between a single locus (SNP) and a single trait. Such studies are not capable of taking advantage of information from multiple correlated traits.

It is well known that many genes may control the variation of more than one trait, a phenomenon known as pleiotropy [2]. Previous studies have shown that not only can a joint analysis of multiple correlated traits increase statistical power for pleiotropic genes but also genes that only affect one of multiple correlated traits [3]. Multivariate analyses can increase power because multivariate records can effectively increase population size relative to univariate records [4]. However, joint analysis of multiple traits is much more time consuming than separate analyses of individual traits due to the complex phenotypic covariance structures among the traits.

Multivariate linear mixed models (mvLMM) are powerful methods used for analysis of multiple correlated traits because of their ability to account for population structures and cryptic relatedness among individuals [5–7]. However, even fitting the mvLMM for a single SNP requires a computational complexity of *O*(*N*^3^), where *N* = *n* × *t* is the total number of phenotypic records for *t* traits and *n* is the sample size. To avoid repeatedly optimizing the variance parameter in testing each SNP, Korte *et al* [7] calculated the covariance matrix only once and only re-estimated a scalar of the covariance structure for every SNP, which has reduced the per-SNP computational time to *O*(*n*^2^). However, the approximate test statistic is associated with a low power. Zhou and Stephens [6] proposed an efficient algorithm using eigen-decomposition of the relatedness matrix, resulting in a time complexity of *O*(*n*^2^) to estimate the variance parameter for each SNP. This eigen-decomposition approach allows exact test for each SNP without compromising the power. So far, all existing methods of multivariate GWAS require complete phenotypic records. Individuals with partial records must be removed from the data prior to the multivariate analysis, resulting in reduction of the sample size and thus low power.

We report here an efficient algorithm and software package named genomic multivariate analysis tool (GMAT), which performs multivariate GWAS including individuals with incomplete phenotypic data. To improve the computational efficiency, we propose an approximate algorithm, which first implements an approximate statistical test with a computational complexity of *O*(*n*) to filter out many non-significant SNPs and then re-test the effects of selected SNP using the exact Wald Chi-square test. Meanwhile, a weighted EM and AI iterative algorithm is applied to improve the computational efficiency in estimating the variance parameters. Simulation studies were conducted to investigate the properties of GMAT. Analyses of data from donkey [8] and yeast [9] showed that GMAT is an efficient method to carry out multivariate GWAS for populations with complete or incomplete phenotypic records.

## 2 Material and methods

### 2.1 GMAT model

To clearly clarify GMAT model, we present a small example with detailed formula derivation in **Supplementary Note**. The linear mixed model for multiple traits can be written as

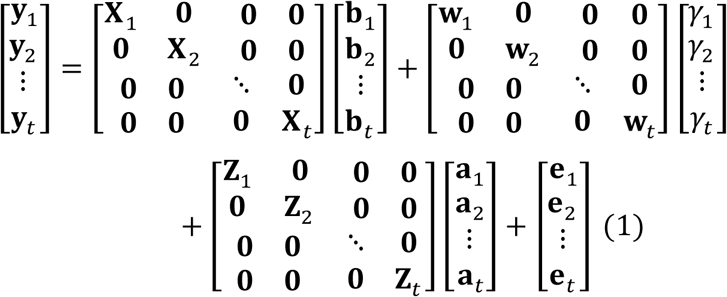

where **y**_*i*_ is a vector of phenotypic values for the *i*th trait; **b**_*i*_ is a vector of fixed effects; **a**_*i*_ is a vector of additive polygenic genetic effects; **e**_*i*_ is a vector of residual errors. **X**_*i*_ and **Z**_*i*_ are the corresponding design matrices for the fixed effects and the polygenic effects. *γ*_*i*_ is the effect of the SNP under study for the *i*th trait and **w**_*i*_ is a vector of SNP genotypes assigned a value of 0, 1 or 2, respectively, for *aa, Aa* and *AA*. Here, the number of phenotypic records may be different for different traits (*i*.*e*., the length of **y**_*i*_ is different for different traits), which means that missing phenotypic records for some traits are allowed. We also refer data with missing phenotypic records to unbalanced data. The number of elements and order in **a**_*i*_ is the same for all traits and the additive genetic effects of individuals without phenotypic records can be estimated from information of related individuals. If we define **w** as a vector of SNP genotypes for all individuals whose order is consistent with **a**_*i*_, we have

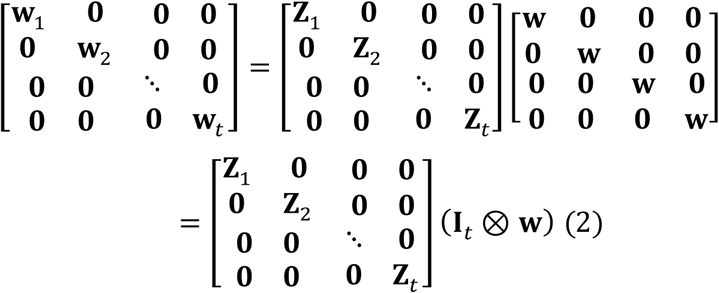

where **I**_*t*_ is a *t* × *t* identity matrix; ⊗ is the Kronecker product. If we define

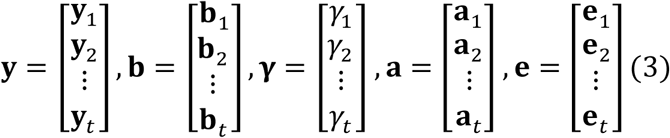

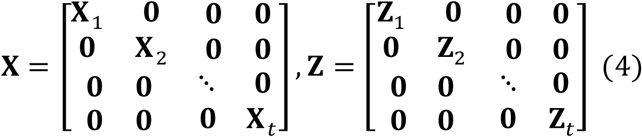

equation (1) can be rewritten as

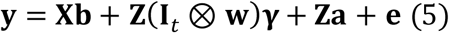

Note that the elements in vector **y** and **a** are in the order that individuals are nested within traits.

The distributions of the random effects are

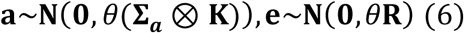

The phenotypic variance is

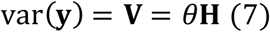

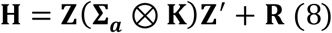

Here, **Σ**_*a*_ is a *t* × *t* covariance matrix for the additive polygenic effects; **R** is a covariance matrix for the residuals; *θ* is the overall scale parameter; **K** is a genomic relationship matrix built from the first method of VanRaden [10]. The variance parameters in the model are estimated with residual maximum likelihood estimation (REML) [11]. The SNP effects of different traits can be estimated from the following equation,

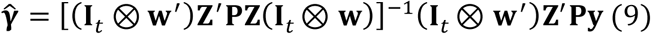

where

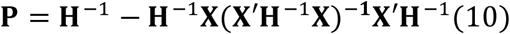

The variance-covariance matrix of the estimated genetic effects is

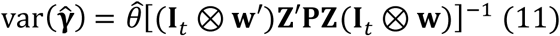

The Wald Chi-square test statistics can be constructed to test the significance of the SNP effects,

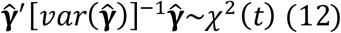

For balanced data that missing phenotypic records are not allowed for all traits (*i*.*e*., the length of **y**_*i*_ is same for different traits), we rearranged the multivariate data so that the traits are nested within individuals. The model is rewritten as

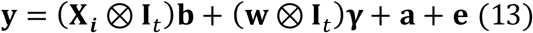

Here, **X**_***i***_ is the same for all traits in the balanced data. Distributions of the random effects are

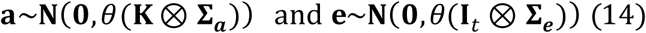

We performed eigen-decomposition for matrix **K (K** = **UDV**′) where **U** is the eigenvector matrix and **D** is a diagonal matrix of eigenvalues. We then multiplied equation (13) by **U**′ ⊗ **I**_*t*_ so that

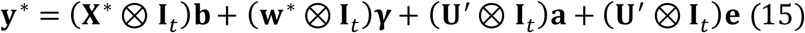

where **y**\* = (**U**′ ⊗ **I**_*t*_)**y, X**\* = **U**′**X**_***i***_, **w**\* = **U**′**w**. The variance matrix of the transformed phenotypes is

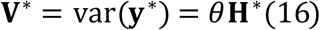

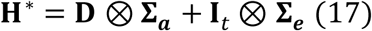

which is a block diagonal matrix. The time complexity to invert this block diagonal matrix is *O*(*nt*^3^). The fact that *t* is usually a much smaller number than *n*, the time complexity for estimating variance components is linear with the number of individuals, which makes it practical to re-estimate the variance parameters for each SNP association analysis. The estimated SNP effects and the corresponding covariance matrix are

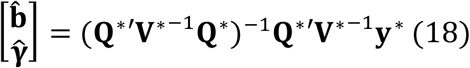

and

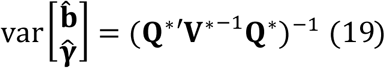

where

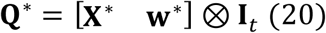

Note that we are estimating **b** and **γ** simultaneously. Since we are only interested in the estimated SNP effects and the variance matrix of the estimated SNP effects, we can take the following alternative formulas to estimate the SNP effects and their covariance matrix,

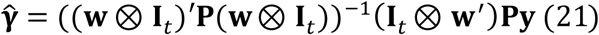

and

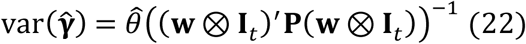

We used a series of efficient algorithms to accelerate the computational efficiency. For estimation of variance components of unbalanced and balanced mvLMM, we combined the expectation & maximization (EM) algorithm with the average information (AI) algorithm to construct a weighted information matrix, which guarantees the variance parameters to converge rapidly within their legal domain [12]. For the unbalanced mvLMM algorithm, it is still time-consuming to estimate the variance components with weighted EM and AI algorithm. To avoid repeatedly estimating the variance parameters for each SNP association analysis, we decided to estimate the covariance structure (**Σ**_***a***_ and **Σ**_***e***_ ) under the null model and only re-estimate the overall scale parameter in testing the genome-wide SNPs using the following equation

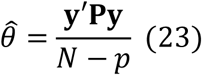

where *N* is the number of phenotypic records; *p* is the rank of design matrix for fixed effects (including SNP effects). For the balanced mvLMM algorithm, we estimated the variances using the transformed mvLMM, equation (15), which allowed estimating variance components under time complexity of *O*(*n*).

With **Z**′**Py** and **Z**′**PZ** being pre-calculated, the time complexity for each SNP association is *O*(*n*^2^). Meanwhile, we developed an efficient algorithm to implement the approximate Wald Chi-squared statistical test, which is described in the following three steps.

**Step 1:** Randomly select *k* (e.g., *k =* 100) SNPs, estimate the effects and calculate the covariance matrix for the estimated SNP effects, using equations (11) and (19) for the unbalanced and balanced data, respectively. We then divide the SNPs into five classes according to their minor allele frequencies (MAF) (0.0-0.1, 0.1-0.2, 0.2-0.3, 0.3-0.4 and 0.4-0.5). Eventually, we estimate the SNP effects and calculate the covariance matrix of the estimated effects for each MAF classes. Finally, we calculate the mean of the covariance matrices across the MAF classes, which is denoted by 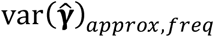. This step has a time complexity of *O*(*kn*^2^).

**Step 2:** Scan the genome-wide SNP effects and calculate the approximate Wald Chi-squared statistical test for each SNP with the following equations. The estimated SNP effects for the unbalanced data are

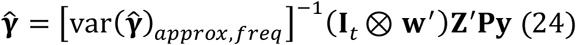

The estimated SNP effects for the balanced data are

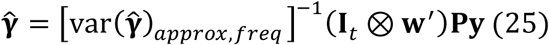

The approximate Wald Chi-squared statistical test is

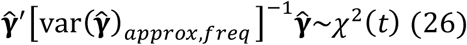

When **Z**′**Py** or **Py** are pre-calculated, the time complexity for this step is simply *O*(*n*).

**Step 3:** Select top significant SNPs (e.g., *P* < 1.0e-3) based on the approximate Wald Chi-squared tests and re-calculate their *P* values using the exact Wald Chi-squared statistical test. The time complexity for this step is *O*(*rn*^2^), where *r* is the number of top significant SNPs.

### 2.2 Simulation

We first investigated the type 1 error of our method and compared it with other existing methods. We calculated the genomic relationship matrix from the original SNPs and then randomly shuffled the SNPs across individuals at each analysis. This random shuffling of the SNPs had purposely destroyed the association of the phenotypes with the scanned SNPs and the linkage disequilibrium (LD) among SNPs. The covariance structure of the original phenotypes induced by the complex cryptic genetic relationship among the individuals remained intact. Under the assumption that random SNPs are not linked to polymorphisms that control the traits of interest, the cumulative *P*-value follows the standard uniform distribution *U*(0, 1). To compare the powers of different models, we used the real genotypes and phenotypes from the mouse data [13] to simulate genetic effects that can recover the original phenotypes. For the mouse data, we used 8312 SNPs from 1642 individuals after removing individuals without genotypes and SNPs with a minor allele frequency (MAF) lower than 0.05 and those that failed the Hardy-Weinberg equilibrium (HWE) test (*P* value < 1.0e-6). We used phenotypes of creatinine, aspartate amino transferase (AST), high density lipoprotein (HDL) and alkaline phosphatase (ALP) to generate the simulated data. The following trait pairs, HDL-ALP, HDL-AST and Creatinine-AST, were analyzed with a one-trait GWAS model and a two-trait GWAS model, where all SNPs with *P* < 0.1 were removed. We then randomly selected 1000 SNPs to act as causal SNPs. For each causal SNP, we specified its effect on the first trait to explain a predetermined proportion of the phenotypic variance (from 0.5% to 5% incremented by 0.5%). Afterward, we specified its pleiotropic effect on the second trait so that the proportion of phenotypic variance explained for the second trait by the SNP equaled 0%, 2% or 4%. To evaluate the performance of different models in dealing with incomplete multivariate data, we randomly removed phenotypes of 25% and 45% individuals for the two traits, respectively. Individuals with missing records for the first trait did not overlap individuals with missing records for the second trait.

### 2.3 Real Datasets

We used two real datasets to evaluate the performance of GMAT, a balanced donkey data [8] and an unbalanced yeast data [9]. The genotypes of the donkey data were obtained with low-coverage whole genome sequence with an average sequencing depth of 3.5x. After a stringent quality control, 573 individuals with 2,298,835 SNPs remained. We analyzed the birth weight (BW) and the weaning weight (WW) in a two-trait linear mixed model. The yeast segregants were genotyped with 28,220 markers (no heterozygous genotype). The traits of cobalt chloride, copper sulfate and raffinose were analyzed with a three-trait model. The average phenotypic values of multiple replications per strain were used as the response variables. After removing strains with missing values, there were 2267, 3109 and 2593 strains for cobalt chloride, copper sulfate and raffinose traits, respectively. The number of strains with at least one recorded trait was 3421.

## 3 Results

We first validated the performance of different models with the simulated data. Here, GMAT-exact and GMAT-approx represent GMAT with the exact and approximate Wald Chi-squared statistical tests, respectively. GMAT-approx2exact represents a scenario that the genome-wide SNPs were scanned with the approximate Wald Chi-squared statistical test and then the top significant SNPs (approximate *P* < 1.0e-3) were re-tested with the exact Wald tests. The quantile-quantile (Q-Q) plots in **Figure 1A** and **Figure 2A** show that the type 1 errors are well controlled by the GMAT-exact and GMAT-approx2exact methods with both the balanced donkey data and the unbalanced yeast data. The GMAT-approx method has slightly inflated *P* values for the donkey data. Comparing the –log_10_(*p*) values obtained from the GMAT-exact and the GEMMA [6] methods, we found that they share exactly the same –log_10_(*p*) values for the donkey data (**Figure 1B**). This is expected as GMAT and GEMMA use the same model when analyzing balanced multiple traits. We compared the –log_10_(p) values between the exact Wald test and the approximate Wald tests (**Figure 1C and Figure 2B**). The two are highly correlated, especially, for the yeast data where the correlations are almost perfect. This may be due to the fact that the MAFs of genome-wide SNPs in the yeast data are very close to 0.5. For the GMAT-approx2exact method, we used the cutoff *P* value of 1.0e-3 to select significant SNPs and then re-calculated these *P* values with the exact test statistics. We found that the *P* values of GMAT-approx2exact and GMAT-exact are the same for all SNPs with *P* values < 1.0e-4 (**Figure 1D and Figure 2C**), which indicated that all the significant SNPs by GMAT-exact can also be located by GMAT-approx2exact after genome-wide Bonferroni correction.

**Figure 1.**
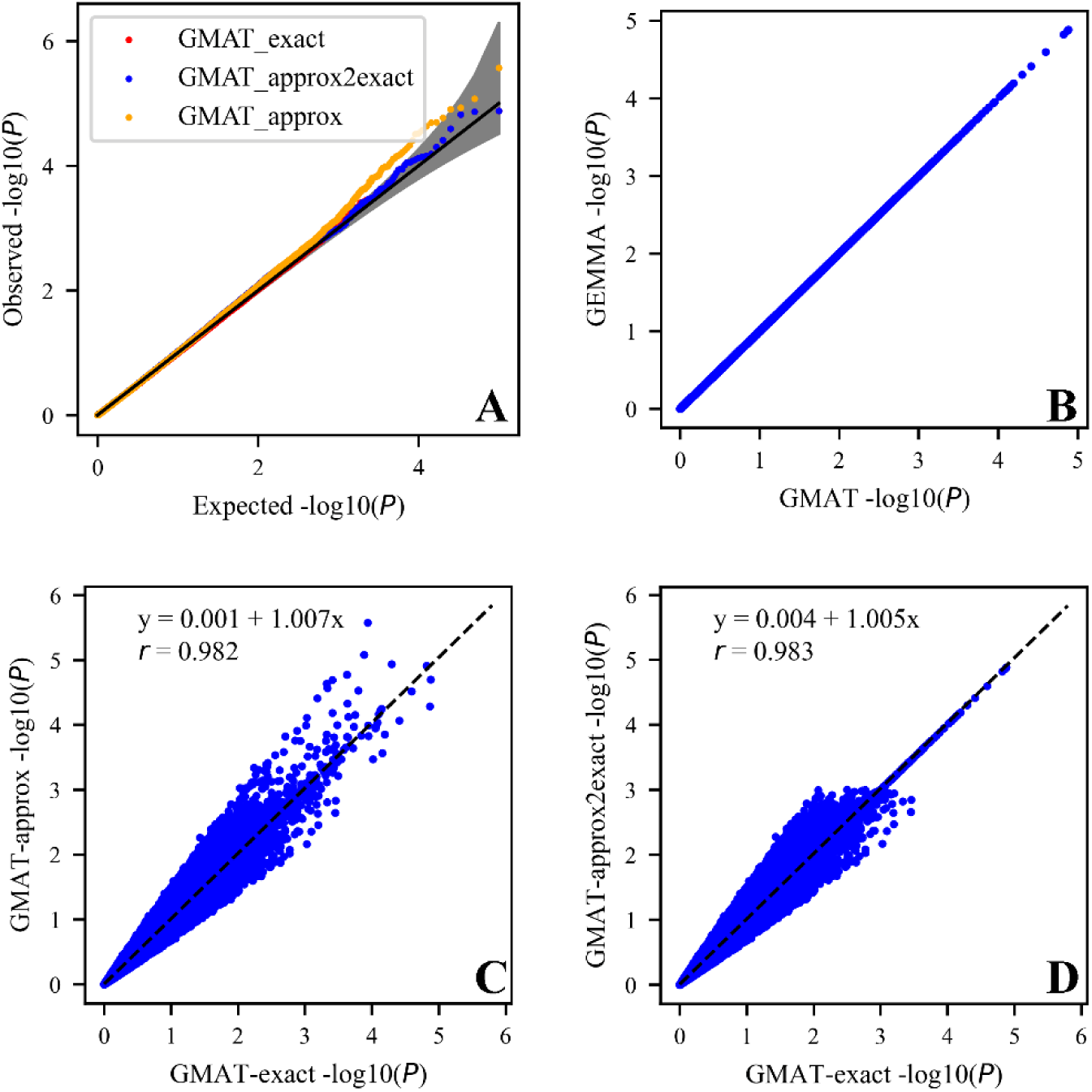
Comparison of the -log_10_(*P*) values for different methods with the balanced donkey data. (A) The quantile-quantile (Q-Q) plots of -log_10_(*P*) for GMAT-exact, GMAT-approx and GMAT-approx2exact; (B) Comparison of -log_10_(*P*) values obtained by GMAT-exact (X axis) and GEMMA (Y axis); (C) Comparison of -log_10_(*P*) values obtained by GMAT-exact (X axis) and GMAT-approx (Y axis); (D) Comparison of -log_10_(*P*) values obtained by GMAT-exact (X axis) and GMAT-approx2exact (Y axis).

**Figure 2.**
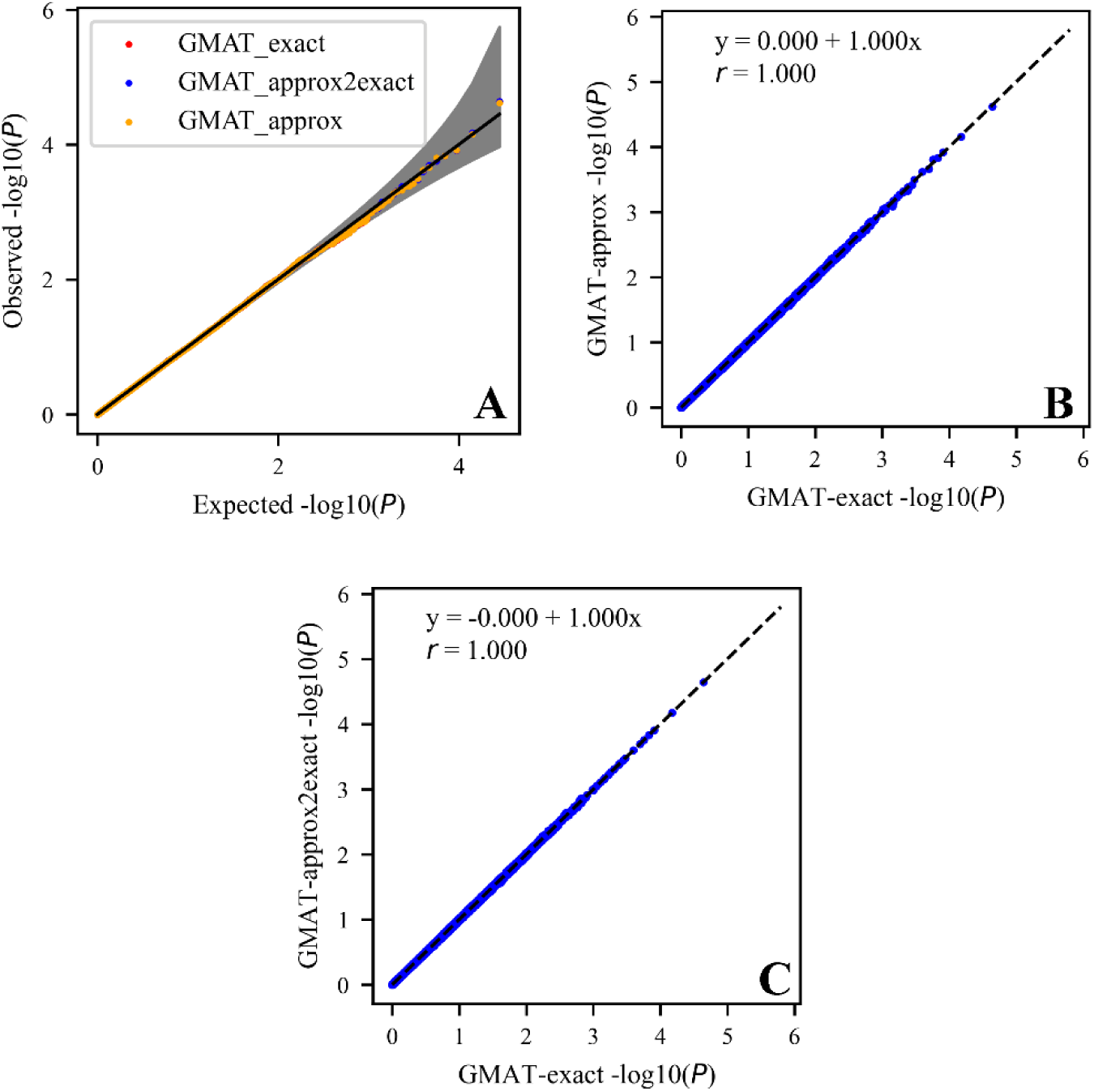
Comparison of the –log10(p) for different methods with the unbalanced yeast data. (A) The quantile-quantile (Q-Q) plots of -log_10_(*P*) for GMAT-exact, GMAT-approx and GMAT-approx2exact; (B) Comparison of -log_10_(*P*) values obtained by GMAT-exact (X axis) and GMAT-approx (Y axis); (C) Comparison of - log_10_(*P*) values obtained by GMAT-exact (X axis) and GMAT-approx2exact (Y axis).

Statistical powers of the simulation experiments for different methods of the genome-wide association studies are shown in **Figure 3**. Here, GMAT-balance represents the method where all individuals with partial records have been removed and individuals with full records were analyzed with the balanced mvLMM algorithm; GMAT-unbalance fits incomplete multivariate data with the unbalanced mvLMM algorithm; GMAT-balance-fill represents a scenario where missing records were imputed with the univariate linear mixed model (uvLMM) and then the data were analyzed with the balanced mvLMM algorithm. In our simulation studies, the statistical powers of GMAT-exact and GMAT-approx2exact are much the same. Therefore, we only show the properties of the GMAT-exact method here. For the simulation experiment of no missing records for two traits (**Figure 3A,D,G**), the GMAT-balance method outperformed the GMAT-unbalance method in terms of higher statistical power. The balanced multivariate data analyzed with the unbalanced mvLMM algorithm has avoided repeated estimation of the variance parameters for each SNP. When we randomly removed a percentage of phenotypic records, the GMAT-balance method has lowered the statistical power. For some simulation experiments with high missing rates, the detection powers of GMAT-balance have been reduced to nearly zero (e.g. **Figure 3C**). If we filled the missing phenotypes with the univariate LMM algorithm and implemented the GWAS with the balanced mvLMM algorithm (GMAT-balance-fill), the detection power has been dramatically improved. The powers of the GMAT-balance-fill method and the GMAT-unbalance method highly depend on the missing rates and the heritability of the traits. For the simulation experiments based on the HDL-ALP trait pair, the imputation accuracies for the missing phenotypes are 0.60 and 0.62 for heritability of 0.48 and 0.55, respectively. Therefore, GMAT-balance-fill surpassed GMAT-unbalance in terms of detection power. When the heritability of traits used for simulation is low (0.09 for Creatinine and 0.14 for AST), the power of GMAT-unbalance is higher than GMAT-balance-fill due to low imputation accuracies for missing phenotypes (0.20 for Creatinine and 0.25 for AST). For simulation experiments based on HDL and AST, GMAT-unbalance performed better than GMAT-balance-fill under the condition of a higher missing rate. We also compared the performance of different methods under different missing rates in the yeast data (**Figure 4**). The results show that GMAT-balance that removes individuals with missing phenotypes has the worst performance. GMAT-balance-fill can detect more QTL regions than GMAT-unbalance when the missing rates are low. However, as the missing rate increased, GMAT-balance-fill failed to detect QTL regions that have been detected by GMAT-unbalance.

**Figure 3.**
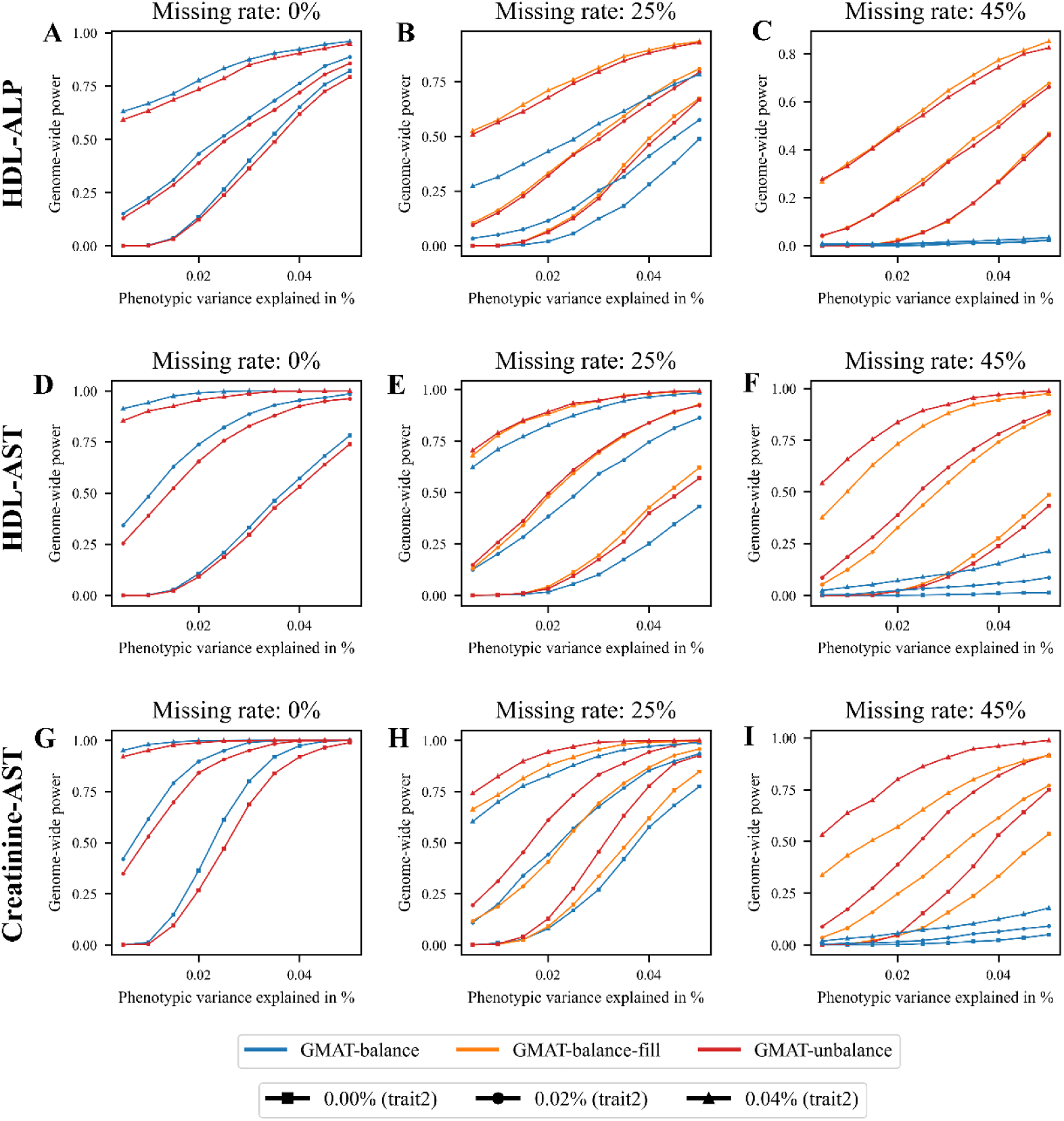
Statistical powers of the mvLMM algorithm for LMM-MGWAS. Panels from the top to the bottom show the result of the simulation experiment based on trait pairs of HDL-ALP, HDL-AST and Creatinine-AST, respectively. The X-axis represents the predetermined proportion of the phenotypic variance explained by the causal SNP for the first trait. The Y-axis represents the genome-wide statistical power after Bonferroni correction. The proportion of the phenotypic variance explained by the causal SNP for the second trait takes three different values: 0% (square), 0.02% (circle) and 0.04% (triangle). Panels from the left to the right show the result of the simulation experiment with missing rates of 0%, 25% and 45%. The GMAT-balance (blue) method is the one that all individuals with missing records have been removed and the data were analyzed using the balanced mvLMM method; The GMAT-unbalance (red) method fits the incomplete multivariate data with unbalanced mvLMM algorithm; The GMAT-balance-fill (orange) method firstly fills the missing phenotypic records with the univariate linear mixed model (uvLMM) and then analyzes the multivariate data with the balanced mvLMM algorithm. For each causal SNP, we specified its effect for the first trait of the trait pair to explain a predetermined proportion of the phenotypic variance.

**Figure 4.**
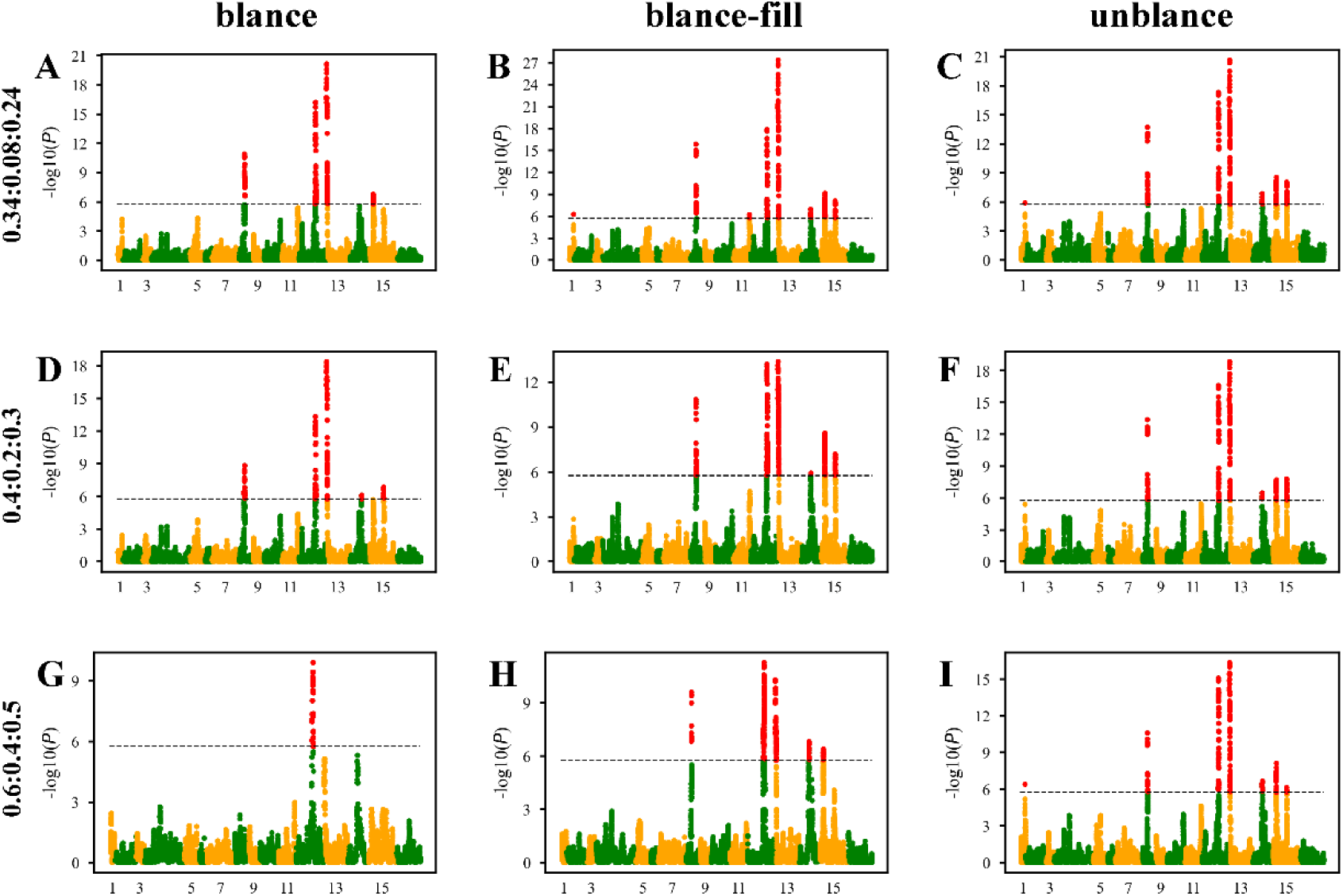
Manhattan plots of the GMAT analysis for the yeast data with three-traits. The three traits are cobalt chloride, copper sulfate and raffinose. The dotted lines indicate the genome-wise significance level (-log_10_(0.05/28,220)) after Bonferroni correction. Missing rates of the three traits are 0.34, 0.08 and 0.24 for the top panels; 0.4, 0.2 and 0.3, for the panels in the middle and 0.6, 0.4 and 0.5 for the panels at the bottom. Panels at the left, the center and the right show the results of GMAT-balance, GMAT-balance-fill and GMAT-unbalance, respectively.

We compared the theoretical time complexity of GMAT and the real computing time of GEMMA and uvLMM (**Table 1**). Both uvLMM and GEMMA have time complexity of *O*(*n*^2^) for the association step of each SNP. For GMAT, the number of random selected SNPs (*s*) can be neglected compared to the total number of SNPs, especially for sequence data. The time complexity of scanning the genome-wide SNPs with the approximate test statistics is linear with the number of individuals (*n*). If we select the top significant SNPs using the *P* < 1.0e-3 criterion, approximately thousands of SNPs are to be re-analyzed with the exact test statistics and the time complexity is *O*(*n*^2^) for each SNP. From the running times of the donkey and yeast data, we can clearly see that GMAT is much faster than GEMMA and uvLMM.

**Table 1.** Theoretical time complexity and actual computational times of different methods.

| Method | Time complexity | Computing time |  |
| --- | --- | --- | --- |
|  |  | Donkey | Yeast |
| uvLMM | $O(mn^2)$ | 34min | 4.85min |
| GEMMA | $O(mn^2)$ | 69.5min | 6min |
| GMAT | $O(sn^2 + mn + rn^2)$ | 1.4min | 20.4s |

Note: Only the performance of the association step was compared in our studies. uvLMM means univariate linear mixed model implemented in gemma software[14]. GEMMA means balanced multivariate linear mixed model implemented in gemma software[6]. All computations were performed on an Intel(R) Xeon(R) Gold 5218 2.3 GHz CPU, where $n$ is the number of individuals; $s$ is the number of randomly selected SNPs; $m$ is the number of SNPs; $r$ is the number of top significant SNPs (approximate $P < 1.0\text{e-}3$ ).

## 4 Discussion

Multivariate GWAS provides an appealing approach to probing the pleiotropic genetic mechanism of complex traits. However, existing multivariate methods require complete multiple phenotypes for each individual and require time cost *O*(*mn*^2^) (where *n* is the number of samples and *m* is the number of SNPs). Here, we developed a far more efficient linear mixed model for multivariate association study (GMAT), which can deal with unbalanced multivariate data and reduce the computational time complexity to a small number of *O*(*mn*). Simulations based on genotypes and phenotypes of actual populations show that GMAT enables an increased association power over methods removing individuals with incomplete records while still properly controls false positives. Applications of the new method to the balanced donkey data and unbalanced yeast data show that the computation efficiency can be increased by dozens of times over existing multivariate linear mixed model association studies. For balanced data, GMAT can achieve the same detection power as GEMMA, but the former is computationally much faster than the latter. This allows GMAT to analyze a large population genotyped with several millions of SNPs. Meanwhile, we developed two methods to deal with unbalanced data, one is to directly fit the data (GMAT-unbalance) and the other is to impute missing phenotypes with the uvLMM algorithm and then analyzed the data with the balanced mvLMM algorithm (GMAT-balance-fill). Performance of the two methods depend on the imputation accuracy of missing records and the missing rates. GMAT-unbalance surpasses GMAT-balance-fill under low filling accuracy and high missing rates. Factors influencing the imputation accuracy may include trait heritability, imputation methods and population size. In real data analyses, it is hard to know the real imputation accuracy a priori and determine which method would have higher powers. The fact that both GMAT-unbalance and GMAT-balance-fill properly control the type 1 error, both methods should be applied and we can use the union set of significant SNPs for the follow up studies.

It is an ideal approach to re-estimate variance parameters at each marker, especially for those with relatively large effects. However, the variance parameters estimation in directly fitting the unbalanced multi-trait data need to invert a dense phenotypic variance matrix, whose time complexity is *O*(*n*^3^). In the analysis of yeast data, the running time for variance parameters per marker is about 6 minutes. A simple extrapolation suggests that it would take more than 118 days to analyze genome-wide SNPs. To improve the computational efficiency, we restarted the estimation with variance parameters from balanced mvLMM, which could reduce the running time by a half. Besides, we fixed the covariance structure (**Σ**_***a***_ and **Σ**_***e***_ ) and only re-estimated the overall scale parameter (*θ*) per SNP, which reduced the time complexity to *O*(*n*^2^). Nevertheless, it still results in power loss for balanced multi-trait data according to the simulation results.

Other comprehensive methods, such as Bayesian and machine learning, can also be applied to improve the imputation accuracy to increase the power of GMAT-balance-fill. However, whether they can perform better than the uvLMM algorithm in prediction will depend on the population size and genetic architecture of traits[15]. In the study, we used the uvLMM algorithm to impute missing phenotypes because of its simplicity and stability in prediction.

## Availability of data and materials

Software is available at https://github.com/chaoning/GMAT. All the datasets analyzed here were from previously published datasets.

## Authors’ contributions

C.N. and Q.Z. conceived and designed the experiments. C.N. and D.W. contributed analytic tools and analysed the data. J.T., H.T., X.F. and S.X. participated in the result interpretation and paper revision. C.N. and D.W. wrote the paper with comments from S.X. All authors read and approved the final manuscript.

## Acknowledgements

Not applicable.

