## Supplementary Note for "An Improved Linear Mixed Model for Multivariate Genome-Wide Association Studies"

**Supplementary Information**

Here is a small example of unbalanced multiple traits. There are three individuals measured with three traits. Missing phenotypic records are expressed as NA.

| ID | sex | snp | trait1 | trait2 | trait3 |
| --- | --- | --- | --- | --- | --- |
| 1 | 0 | 0 | NA | 8.4 | NA |
| 2 | 1 | 1 | 4.6 | NA | 10.9 |
| 3 | 1 | 2 | 8 | 9.7 | NA |

The linear mixed model for the *i*th trait can be written as

$$\mathbf{y}_{i}=\mathbf{X}_{i}\mathbf{b}_{i}+\mathbf{w}_{i}\gamma_{i}+\mathbf{Z}_{i}\mathbf{a}_{i}+\mathbf{e}_{i} (1)$$

where $\mathbf{y}_{i}$ is a vector of phenotypic values for the $i$th trait; $\mathbf{b}_{i}$ is a vector of fixed effects (population mean and sex); $\mathbf{a}_{i}$ is a vector of additive polygenic genetic effects; $\mathbf{e}_{i}$ is a vector of residual errors. $\mathbf{X}_{i}$ and $\mathbf{Z}_{i}$ are the corresponding design matrices for the fixed effects and the polygenic effects. $\gamma_{i}$ is the effect of the SNP under study for the $i$th trait and $\mathbf{w}_{i}$ is a vector of SNP genotypes assigned a value of 0, 1 or 2, respectively, for *aa*, *Aa* and *AA*. We define $\mathbf{w}$ as a vector of SNP genotypes for all individuals and $\mathbf{e}_{i}^{*}$ as a vector of pseudo residual errors including missing phenotypic records. Both have order consistent with $\mathbf{a}_{i}$. Then,

$$\mathbf{w}_{i}=\mathbf{Z}_{i}\mathbf{W}(2)$$

$$\mathbf{e}_{i}=\mathbf{Z}_{i}\mathbf{e}_{i}^{*} (3)$$

For the three traits in the example, we can write as

$$\left[ \begin{matrix} y_{12} \\ y_{13} \end{matrix} \right]=\left[ \begin{matrix} 1 & 1 \\ 1 & 1 \end{matrix} \right]\left[ \begin{matrix} \mu_{1} \\ s_{1} \end{matrix} \right]+\left[ \begin{matrix} 1 \\ 2 \end{matrix} \right]\gamma_{1}+\left[ \begin{matrix} 0 & 1 & 0 \\ 0 & 0 & 1 \end{matrix} \right]\left[ \begin{matrix} a_{11} \\ a_{12} \\ a_{13} \end{matrix} \right]+\left[ \begin{matrix} e_{12} \\ e_{13} \end{matrix} \right]=\left[ \begin{matrix} 1 & 1 \\ 1 & 1 \end{matrix} \right]\left[ \begin{matrix} \mu_{1} \\ s_{1} \end{matrix} \right]+\left[ \begin{matrix} 0 & 1 & 0 \\ 0 & 0 & 1 \end{matrix} \right]\left[ \begin{matrix} 0 \\ 1 \\ 2 \end{matrix} \right]\gamma_{1}+\left[ \begin{matrix} 0 & 1 & 0 \\ 0 & 0 & 1 \end{matrix} \right]\left[ \begin{matrix} a_{11} \\ a_{12} \\ a_{13} \end{matrix} \right]+\left[ \begin{matrix} 0 & 1 & 0 \\ 0 & 0 & 1 \end{matrix} \right]\left[ \begin{matrix} e_{11}^{*} \\ e_{12} \\ e_{13} \end{matrix} \right] (4)$$

$$\left[ \begin{matrix} y_{21} \\ y_{23} \end{matrix} \right]=\left[ \begin{matrix} 1 & 0 \\ 1 & 1 \end{matrix} \right]\left[ \begin{matrix} \mu_{2} \\ s_{2} \end{matrix} \right]+\left[ \begin{matrix} 0 \\ 2 \end{matrix} \right]\gamma_{2}+\left[ \begin{matrix} 1 & 0 & 0 \\ 0 & 0 & 1 \end{matrix} \right]\left[ \begin{matrix} a_{21} \\ a_{22} \\ a_{23} \end{matrix} \right]+\left[ \begin{matrix} e_{21} \\ e_{23} \end{matrix} \right]=\left[ \begin{matrix} 1 & 0 \\ 1 & 1 \end{matrix} \right]\left[ \begin{matrix} \mu_{2} \\ s_{2} \end{matrix} \right]+\left[ \begin{matrix} 1 & 0 & 0 \\ 0 & 0 & 1 \end{matrix} \right]\left[ \begin{matrix} 0 \\ 1 \\ 2 \end{matrix} \right]\gamma_{2}+\left[ \begin{matrix} 1 & 0 & 0 \\ 0 & 0 & 1 \end{matrix} \right]\left[ \begin{matrix} a_{21} \\ a_{22} \\ a_{23} \end{matrix} \right]+\left[ \begin{matrix} 1 & 0 & 0 \\ 0 & 0 & 1 \end{matrix} \right]\left[ \begin{matrix} e_{21} \\ e_{22}^{*} \\ e_{23} \end{matrix} \right] (5)$$

$$\left[ y_{32} \right]=\left[ \begin{matrix} 1 & 1 \end{matrix} \right]\left[ \begin{matrix} \mu_{3} \\ s_{3} \end{matrix} \right]+\left[ 1 \right]\gamma_{3}+\left[ \begin{matrix} 0 & 1 & 0 \end{matrix} \right]\left[ \begin{matrix} a_{21} \\ a_{22} \\ a_{23} \end{matrix} \right]+\left[ e_{32} \right]=\left[ \begin{matrix} 1 & 1 \end{matrix} \right]\left[ \begin{matrix} \mu_{3} \\ s_{3} \end{matrix} \right]+\left[ \begin{matrix} 0 & 1 & 0 \end{matrix} \right]\left[ \begin{matrix} 0 \\ 1 \\ 2 \end{matrix} \right]\gamma_{3}+\left[ \begin{matrix} 0 & 1 & 0 \end{matrix} \right]\left[ \begin{matrix} a_{21} \\ a_{22} \\ a_{23} \end{matrix} \right]+\left[ \begin{matrix} 0 & 1 & 0 \end{matrix} \right]\left[ \begin{matrix} e_{31}^{*} \\ e_{32} \\ e_{33}^{*} \end{matrix} \right] (6)$$

Here, $y_{ij}$ is the phenotypic value for the $i$th trait of individual $j$; $a_{ij}$ is the additive polygenic genetic effect; $e_{ij}$ is the residual error, $e_{ij}^{*}$ is the pseudo residual error for one missing phenotypic record.

The linear mixed model for three traits can be written as

$$\left[ \begin{matrix} \mathbf{y}_{1} \\ \mathbf{y}_{2} \\ \mathbf{y}_{3} \end{matrix} \right]=\left[ \begin{matrix} \mathbf{X}_{1} & \mathbf{0} & \mathbf{0} \\ \mathbf{0} & \mathbf{X}_{2} & \mathbf{0} \\ \mathbf{0} & \mathbf{0} & \mathbf{X}_{3} \end{matrix} \right]\left[ \begin{matrix} \mathbf{b}_{1} \\ \mathbf{b}_{2} \\ \mathbf{b}_{3} \end{matrix} \right]+\left[ \begin{matrix} \mathbf{Z}_{1}\mathbf{W} & \mathbf{0} & \mathbf{0} \\ \mathbf{0} & \mathbf{Z}_{2}\mathbf{W} & \mathbf{0} \\ \mathbf{0} & \mathbf{0} & \mathbf{Z}_{3}\mathbf{W} \end{matrix} \right]\left[ \begin{matrix} \gamma_{1} \\ \gamma_{2} \\ \gamma_{3} \end{matrix} \right]+\left[ \begin{matrix} \mathbf{Z}_{1} & \mathbf{0} & \mathbf{0} \\ \mathbf{0} & \mathbf{Z}_{2} & \mathbf{0} \\ \mathbf{0} & \mathbf{0} & \mathbf{Z}_{3} \end{matrix} \right]\left[ \begin{matrix} \mathbf{a}_{1} \\ \mathbf{a}_{2} \\ \mathbf{a}_{3} \end{matrix} \right]+\left[ \begin{matrix} \mathbf{Z}_{1}\mathbf{e}_{1}^{*} \\ \mathbf{Z}_{2}\mathbf{e}_{2}^{*} \\ \mathbf{Z}_{3}\mathbf{e}_{3}^{*} \end{matrix} \right](7)$$

We have

$$\left[ \begin{matrix} \mathbf{Z}_{1}\mathbf{W} & \mathbf{0} & \mathbf{0} \\ \mathbf{0} & \mathbf{Z}_{2}\mathbf{W} & \mathbf{0} \\ \mathbf{0} & \mathbf{0} & \mathbf{Z}_{3}\mathbf{W} \end{matrix} \right]=\left[ \begin{matrix} \mathbf{Z}_{1} & \mathbf{0} & \mathbf{0} \\ \mathbf{0} & \mathbf{Z}_{2} & \mathbf{0} \\ \mathbf{0} & \mathbf{0} & \mathbf{Z}_{3} \end{matrix} \right]\left[ \begin{matrix} \mathbf{W} & \mathbf{0} & \mathbf{0} \\ \mathbf{0} & \mathbf{W} & \mathbf{0} \\ \mathbf{0} & \mathbf{0} & \mathbf{W} \end{matrix} \right]=\left[ \begin{matrix} \mathbf{Z}_{1} & \mathbf{0} & \mathbf{0} \\ \mathbf{0} & \mathbf{Z}_{2} & \mathbf{0} \\ \mathbf{0} & \mathbf{0} & \mathbf{Z}_{3} \end{matrix} \right]\left( \mathbf{I}_{3}\otimes\mathbf{w} \right) (8)$$

$$\left[ \begin{matrix} \mathbf{Z}_{1}\mathbf{e}_{1}^{*} \\ \mathbf{Z}_{2}\mathbf{e}_{2}^{*} \\ \mathbf{Z}_{3}\mathbf{e}_{3}^{*} \end{matrix} \right]=\left[ \begin{matrix} \mathbf{Z}_{1} & \mathbf{0} & \mathbf{0} \\ \mathbf{0} & \mathbf{Z}_{2} & \mathbf{0} \\ \mathbf{0} & \mathbf{0} & \mathbf{Z}_{3} \end{matrix} \right]\left[ \begin{matrix} \mathbf{e}_{1}^{*} \\ \mathbf{e}_{2}^{*} \\ \mathbf{e}_{3}^{*} \end{matrix} \right] (9)$$

Substitute them into the equation (7), we have

$$\left[ \begin{matrix} \mathbf{y}_{1} \\ \mathbf{y}_{2} \\ \mathbf{y}_{3} \end{matrix} \right]=\left[ \begin{matrix} \mathbf{X}_{1} & \mathbf{0} & \mathbf{0} \\ \mathbf{0} & \mathbf{X}_{2} & \mathbf{0} \\ \mathbf{0} & \mathbf{0} & \mathbf{X}_{3} \end{matrix} \right]\left[ \begin{matrix} \mathbf{b}_{1} \\ \mathbf{b}_{2} \\ \mathbf{b}_{3} \end{matrix} \right]+\left[ \begin{matrix} \mathbf{Z}_{1} & \mathbf{0} & \mathbf{0} \\ \mathbf{0} & \mathbf{Z}_{2} & \mathbf{0} \\ \mathbf{0} & \mathbf{0} & \mathbf{Z}_{3} \end{matrix} \right]\left( \mathbf{I}_{3}\otimes\mathbf{w} \right)\left[ \begin{matrix} \gamma_{1} \\ \gamma_{2} \\ \gamma_{3} \end{matrix} \right]+\left[ \begin{matrix} \mathbf{Z}_{1} & \mathbf{0} & \mathbf{0} \\ \mathbf{0} & \mathbf{Z}_{2} & \mathbf{0} \\ \mathbf{0} & \mathbf{0} & \mathbf{Z}_{3} \end{matrix} \right]\left[ \begin{matrix} \mathbf{a}_{1} \\ \mathbf{a}_{2} \\ \mathbf{a}_{3} \end{matrix} \right]+\left[ \begin{matrix} \mathbf{Z}_{1} & \mathbf{0} & \mathbf{0} \\ \mathbf{0} & \mathbf{Z}_{2} & \mathbf{0} \\ \mathbf{0} & \mathbf{0} & \mathbf{Z}_{3} \end{matrix} \right]\left[ \begin{matrix} \mathbf{e}_{1}^{*} \\ \mathbf{e}_{2}^{*} \\ \mathbf{e}_{3}^{*} \end{matrix} \right](10)$$

where $\mathbf{I}_{3}$ is a $3\times3$ identity matrix; $\otimes$ is the Kronecker product.

The distributions of the random effects are

$$\left[ \begin{matrix} \mathbf{a}_{1} \\ \mathbf{a}_{2} \\ \mathbf{a}_{3} \end{matrix} \right]\sim N\left( \left[ \begin{matrix} \mathbf{0} \\ \mathbf{0} \\ \mathbf{0} \end{matrix} \right], \left[ \begin{matrix} \mathbf{K}\sigma_{a,11}^{2} & \mathbf{K}\sigma_{a,12}^{2} & \mathbf{K}\sigma_{a,13}^{2} \\ \mathbf{K}\sigma_{a,21}^{2} & \mathbf{K}\sigma_{a,22}^{2} & \mathbf{K}\sigma_{a,23}^{2} \\ \mathbf{K}\sigma_{a,31}^{2} & \mathbf{K}\sigma_{a,32}^{2} & \mathbf{K}\sigma_{a,33}^{2} \end{matrix} \right] \right)=N\left( \mathbf{0}, \boldsymbol{\Sigma}_{\boldsymbol{a}}\boldsymbol{\otimes}\mathbf{K} \right) (11)$$

$$\left[ \begin{matrix} \mathbf{e}_{1}^{*} \\ \mathbf{e}_{2}^{*} \\ \mathbf{e}_{3}^{*} \end{matrix} \right]\sim N\left( \left[ \begin{matrix} \mathbf{0} \\ \mathbf{0} \\ \mathbf{0} \end{matrix} \right], \left[ \begin{matrix} \mathbf{I}\sigma_{e,11}^{2} & \mathbf{I}\sigma_{e,12}^{2} & \mathbf{I}\sigma_{e,13}^{2} \\ \mathbf{I}\sigma_{e,21}^{2} & \mathbf{I}\sigma_{e,22}^{2} & \mathbf{I}\sigma_{e,23}^{2} \\ \mathbf{I}\sigma_{e,31}^{2} & \mathbf{I}\sigma_{e,32}^{2} & \mathbf{I}\sigma_{e,33}^{2} \end{matrix} \right] \right)=N\left( \mathbf{0}, \boldsymbol{\Sigma}_{\boldsymbol{e}}\boldsymbol{\otimes}\mathbf{I} \right) (12)$$

Here, $\boldsymbol{\Sigma}_{a}$ is a $3\times3$ covariance matrix for the additive polygenic effects; $\boldsymbol{\Sigma}_{e}$ is a covariance matrix for the residuals; $\mathbf{K}$ is a genomic relationship matrix.

Here is a small example of balanced multiple traits.

| ID | sex | snp | trait1 | trait2 | trait3 |
| --- | --- | --- | --- | --- | --- |
| 1 | 0 | 0 | 5.2 | 8.4 | 11.2 |
| 2 | 1 | 1 | 4.6 | 7.8 | 10.9 |
| 3 | 1 | 2 | 8 | 9.7 | 9.6 |

The linear mixed model for the *i*th trait can be written as

$$\mathbf{y}_{i}=\mathbf{X}_{i}\mathbf{b}_{i}+\mathbf{w}\gamma_{i}+\mathbf{Z}_{i}\mathbf{a}_{i}+\mathbf{e}_{i} (13)$$

Here, $\mathbf{X}_{i}$ is same for all traits (define as **X**); $\mathbf{Z}_{i}$ is an identity matrix. For the three traits in the example, we can write as

$$\left[ \begin{matrix} y_{11} \\ y_{12} \\ y_{13} \end{matrix} \right]=\left[ \begin{matrix} 1 & 0 \\ 1 & 1 \\ 1 & 1 \end{matrix} \right]\left[ \begin{matrix} \mu_{1} \\ s_{1} \end{matrix} \right]+\left[ \begin{matrix} a_{11} \\ a_{12} \\ a_{13} \end{matrix} \right]+\left[ \begin{matrix} e_{11} \\ e_{12} \\ e_{13} \end{matrix} \right] (14)$$

$$\left[ \begin{matrix} y_{21} \\ y_{22} \\ y_{23} \end{matrix} \right]=\left[ \begin{matrix} 1 & 0 \\ 1 & 1 \\ 1 & 1 \end{matrix} \right]\left[ \begin{matrix} \mu_{2} \\ s_{2} \end{matrix} \right]+\left[ \begin{matrix} a_{21} \\ a_{22} \\ a_{23} \end{matrix} \right]+\left[ \begin{matrix} e_{21} \\ e_{22} \\ e_{23} \end{matrix} \right] (15)$$

$$\left[ \begin{matrix} y_{31} \\ y_{32} \\ y_{33} \end{matrix} \right]=\left[ \begin{matrix} 1 & 0 \\ 1 & 1 \\ 1 & 1 \end{matrix} \right]\left[ \begin{matrix} \mu_{3} \\ s_{3} \end{matrix} \right]+\left[ \begin{matrix} a_{31} \\ a_{32} \\ a_{33} \end{matrix} \right]+\left[ \begin{matrix} e_{31} \\ e_{32} \\ e_{33} \end{matrix} \right] (16)$$

The linear mixed model for three traits can be written as

$$\left[ \begin{matrix} \mathbf{y}_{1} \\ \mathbf{y}_{2} \\ \mathbf{y}_{3} \end{matrix} \right]=\left[ \begin{matrix} \mathbf{X} & \mathbf{0} & \mathbf{0} \\ \mathbf{0} & \mathbf{X} & \mathbf{0} \\ \mathbf{0} & \mathbf{0} & \mathbf{X} \end{matrix} \right]\left[ \begin{matrix} \mathbf{b}_{1} \\ \mathbf{b}_{2} \\ \mathbf{b}_{3} \end{matrix} \right]+\left[ \begin{matrix} \mathbf{w} & \mathbf{0} & \mathbf{0} \\ \mathbf{0} & \mathbf{w} & \mathbf{0} \\ \mathbf{0} & \mathbf{0} & \mathbf{w} \end{matrix} \right]\left[ \begin{matrix} \gamma_{1} \\ \gamma_{2} \\ \gamma_{3} \end{matrix} \right]+\left[ \begin{matrix} \mathbf{a}_{1} \\ \mathbf{a}_{2} \\ \mathbf{a}_{3} \end{matrix} \right]+\left[ \begin{matrix} \mathbf{e}_{1} \\ \mathbf{e}_{2} \\ \mathbf{e}_{3} \end{matrix} \right] (17)$$

$$\left[ \begin{matrix} \mathbf{y}_{1} \\ \mathbf{y}_{2} \\ \mathbf{y}_{3} \end{matrix} \right]=(\mathbf{I}_{3}\otimes\mathbf{X})\left[ \begin{matrix} \mathbf{b}_{1} \\ \mathbf{b}_{2} \\ \mathbf{b}_{3} \end{matrix} \right]+(\mathbf{I}_{3}\otimes\mathbf{w})\left[ \begin{matrix} \gamma_{1} \\ \gamma_{2} \\ \gamma_{3} \end{matrix} \right]+\left[ \begin{matrix} \mathbf{a}_{1} \\ \mathbf{a}_{2} \\ \mathbf{a}_{3} \end{matrix} \right]+\left[ \begin{matrix} \mathbf{e}_{1} \\ \mathbf{e}_{2} \\ \mathbf{e}_{3} \end{matrix} \right] (18)$$

We have

$$\left[ \begin{matrix} \mathbf{y}_{1} \\ \mathbf{y}_{2} \\ \mathbf{y}_{3} \end{matrix} \right]=vec\left( \left[ \begin{matrix} {\mathbf{y}^{\mathbf{'}}}_{1} \\ {\mathbf{y}^{\mathbf{'}}}_{2} \\ {\mathbf{y}^{\mathbf{'}}}_{3} \end{matrix} \right]^{'} \right), \left[ \begin{matrix} \mathbf{a}_{1} \\ \mathbf{a}_{2} \\ \mathbf{a}_{3} \end{matrix} \right]=\mathrm{vec}\left( \left[ \begin{matrix} {\mathbf{a}^{\mathbf{'}}}_{1} \\ {\mathbf{a}^{\mathbf{'}}}_{2} \\ {\mathbf{a}^{\mathbf{'}}}_{3} \end{matrix} \right]^{'} \right), \left[ \begin{matrix} \mathbf{e}_{1} \\ \mathbf{e}_{2} \\ \mathbf{e}_{3} \end{matrix} \right]=\mathrm{vec}\left( \left[ \begin{matrix} {\mathbf{e}^{\mathbf{'}}}_{1} \\ {\mathbf{e}^{\mathbf{'}}}_{2} \\ {\mathbf{e}^{\mathbf{'}}}_{3} \end{matrix} \right]^{'} \right) (19)$$

A permutation matrix, denoted $\mathbf{P}_{nt}$, is defined by,

$$\mathbf{P}_{nt}\mathrm{vec}\left( \left[ \begin{matrix} {\mathbf{y}^{\mathbf{'}}}_{1} \\ {\mathbf{y}^{\mathbf{'}}}_{2} \\ {\mathbf{y}^{\mathbf{'}}}_{3} \end{matrix} \right]^{'} \right)=\mathrm{vec}\left( \left[ \begin{matrix} {\mathbf{y}^{\mathbf{'}}}_{1} \\ {\mathbf{y}^{\mathbf{'}}}_{2} \\ {\mathbf{y}^{\mathbf{'}}}_{3} \end{matrix} \right] \right) (20)$$

We can obtain that

$$\mathbf{P}_{nt}\mathrm{vec}\left( \left[ \begin{matrix} {\mathbf{a}^{\mathbf{'}}}_{1} \\ {\mathbf{a}^{\mathbf{'}}}_{2} \\ {\mathbf{a}^{\mathbf{'}}}_{3} \end{matrix} \right]^{'} \right)=\mathrm{vec}\left( \left[ \begin{matrix} {\mathbf{a}^{\mathbf{'}}}_{1} \\ {\mathbf{a}^{\mathbf{'}}}_{2} \\ {\mathbf{a}^{\mathbf{'}}}_{3} \end{matrix} \right] \right) (21)$$

$$\mathbf{P}_{nt}\mathrm{vec}\left( \left[ \begin{matrix} {\mathbf{e}^{\mathbf{'}}}_{1} \\ {\mathbf{e}^{\mathbf{'}}}_{2} \\ {\mathbf{e}^{\mathbf{'}}}_{3} \end{matrix} \right]^{'} \right)=\mathrm{vec}\left( \left[ \begin{matrix} {\mathbf{e}^{\mathbf{'}}}_{1} \\ {\mathbf{e}^{\mathbf{'}}}_{2} \\ {\mathbf{e}^{\mathbf{'}}}_{3} \end{matrix} \right] \right) (22)$$

$$\mathbf{P}_{nt}\left( \mathbf{I}_{3}\otimes\mathbf{X} \right)=\left( \mathbf{X}{\otimes\mathbf{I}}_{3} \right) (23)$$

$$\mathbf{P}_{nt}(\mathbf{I}_{3}\otimes\mathbf{w})=\left( \mathbf{w}{\otimes\mathbf{I}}_{3} \right) (24)$$

We then multiplied equation (18) by $\mathbf{P}_{nt}$ so that

$$\mathrm{vec}\left( \left[ \begin{matrix} {\mathbf{y}^{\mathbf{'}}}_{1} \\ {\mathbf{y}^{\mathbf{'}}}_{2} \\ {\mathbf{y}^{\mathbf{'}}}_{3} \end{matrix} \right] \right)=\left( \mathbf{X}{\otimes\mathbf{I}}_{3} \right)\left[ \begin{matrix} \mathbf{b}_{1} \\ \mathbf{b}_{2} \\ \mathbf{b}_{3} \end{matrix} \right]+\left( \mathbf{w}{\otimes\mathbf{I}}_{3} \right)\left[ \begin{matrix} \gamma_{1} \\ \gamma_{2} \\ \gamma_{3} \end{matrix} \right]+\mathrm{vec}\left( \left[ \begin{matrix} {\mathbf{a}^{\mathbf{'}}}_{1} \\ {\mathbf{a}^{\mathbf{'}}}_{2} \\ {\mathbf{a}^{\mathbf{'}}}_{3} \end{matrix} \right] \right)+\mathrm{vec}\left( \left[ \begin{matrix} {\mathbf{e}^{\mathbf{'}}}_{1} \\ {\mathbf{e}^{\mathbf{'}}}_{2} \\ {\mathbf{e}^{\mathbf{'}}}_{3} \end{matrix} \right] \right) (25)$$

In this way, multivariate data was rearranged so that the traits are nested within individuals.

Distributions of the random effects are

$$\mathrm{var}\left\{ \mathrm{vec}\left( \left[ \begin{matrix} {\mathbf{a}^{\mathbf{'}}}_{1} \\ {\mathbf{a}^{\mathbf{'}}}_{2} \\ {\mathbf{a}^{\mathbf{'}}}_{3} \end{matrix} \right] \right) \right\}=\mathrm{var}\left\{ \mathbf{P}_{nt}\left[ \begin{matrix} \mathbf{a}_{1} \\ \mathbf{a}_{2} \\ \mathbf{a}_{3} \end{matrix} \right] \right\}=\mathbf{P}_{nt}\left( \boldsymbol{\Sigma}_{\boldsymbol{a}}\boldsymbol{\otimes}\mathbf{K} \right){\mathbf{P}^{\mathbf{'}}}_{nt}=\boldsymbol{K\otimes}\boldsymbol{\Sigma}_{\boldsymbol{a}} (26)$$

$$\mathrm{var}\left\{ \mathrm{vec}\left( \left[ \begin{matrix} {\mathbf{e}^{\mathbf{'}}}_{1} \\ {\mathbf{e}^{\mathbf{'}}}_{2} \\ {\mathbf{e}^{\mathbf{'}}}_{3} \end{matrix} \right] \right) \right\}=\mathrm{var}\left\{ \mathbf{P}_{nt}\left[ \begin{matrix} \mathbf{e}_{1} \\ \mathbf{e}_{2} \\ \mathbf{e}_{3} \end{matrix} \right] \right\}=\mathbf{P}_{nt}\left( \boldsymbol{\Sigma}_{\boldsymbol{e}}\boldsymbol{\otimes}\mathbf{I} \right){\mathbf{P}^{\mathbf{'}}}_{nt}=\mathbf{I}_{t}\boldsymbol{\otimes}\boldsymbol{\Sigma}_{\boldsymbol{e}} (27)$$
